# Gradients of connectivity distance in the cerebral cortex of the macaque monkey

**DOI:** 10.1101/467860

**Authors:** Sabine Oligschläger, Ting Xu, Blazej M. Baczkowski, Marcel Falkiewicz, Arnaud Falchier, Gary Linn, Daniel S. Margulies

## Abstract

Cortical connectivity conforms to a series of organizing principles that are common across species. Spatial proximity, similar cortical type, and similar connectional profile all constitute factors for determining the connectivity between cortical regions. We previously demonstrated another principle of connectivity that is closely related to the spatial layout of the cerebral cortex. Using functional connectivity from resting-state fMRI in the human cortex, we found that the further a region is located from primary cortex, the more distant are its functional connections with other areas of the cortex. However, it remains unknown whether this relationship between cortical layout and connectivity extends to other primate species. Here, we investigated this relationship using both resting-state functional connectivity as well as gold-standard tract-tracing connectivity in the macaque monkey cortex. For both measures of connectivity, we found a gradient of connectivity distance extending between primary and frontoparietal regions. As in the human cortex, the further a region is located from primary areas, the stronger its connections to distant portions of the cortex, with connectivity distance highest in frontal and parietal regions. The similarity between the human and macaque findings provide evidence for a phylogenetically conserved relationship between the spatial layout of cortical areas and connectivity.

## Introduction

Cortical connectivity conforms to a series of organizing principles. Regions tend to be more interconnected when they lie close to each other (Betzel et al., 2016; Beul, Grant, & Hilgetag, 2015; Ercsey-Ravasz et al., 2013; Goulas, Uylings, & Hilgetag, 2017; Kaiser & Hilgetag, 2006; Roberts et al., 2016; Rubinov, Ypma, Watson, & Bullmore, 2015; Vértes et al., 2012), when they feature similar cortical microstructure (Barbas, 2015; Beul, Barbas, & Hilgetag, 2017; Beul et al., 2015; Goulas et al., 2017, 2016; Huntenburg et al., 2017; Pandya, Petrides, & Cipolloni, 2015; Pandya & Sanides, 1973; Pandya & Yeterian, 1985), and when they share common connections to other cortical regions (Betzel et al., 2016; Costa, Kaiser, & Hilgetag, 2007; Song, Kennedy, & Wang, 2014; Vértes et al., 2012). These principles are shared across a variety of species such as the mouse, cat, macaque, and human, demonstrating phylogenetically conserved patterns of connectivity in the mammalian brain.

In the human cerebral cortex, we previously described a principle of cortical organization that relates patterns of connectivity to the spatial layout of cortical regions (Oligschläger et al., 2017). Using resting-state fMRI, we revealed a gradient of connectivity distance that progresses from primary sensory-motor areas to higher-order association cortex. Specifically, the further a region is located from primary cortex, the more distant are its functional connections with other areas of the cortex. Connectivity distance peaked in regions of the default mode network (DMN) — regions that underwent disproportionate expansion during primate phylogeny (Hill et al., 2010) and systematically occupy locations at maximal distance from primary cortex (Margulies et al., 2016).

Buckner and Krienen addressed the spatial distribution of connectivity distance in their ‘tethering hypothesis’ (2013) and attributed it to the cortical expansion that occurred during primate phylogeny. Expansion of the cortical sheet is presumed to have caused increasingly larger proportions of cortex to lie distant from developmental constraints that differentiate primary cortex (Buckner & Krienen, 2013; Rosa & Tweedale, 2005). While this link to expansion suggests a phylogenetically conserved relationship between cortical layout and connectivity in other primate species, a direct assessment in nonhuman primates is lacking. We hypothesize that connectivity distance follows a gradient that is anchored in regions of primary cortex in the non-human primate as well. It is thus our goal in the present study to examine the relationship between cortical layout and connectivity in a nonhuman primate species.

In light of our prior finding in the human cerebral cortex, here we investigated whether the relationship between connectivity and cortical layout was consistent in both functional connectivity and gold-standard tract-tracing connectivity data in the macaque monkey. Functional connectivity was calculated based on resting-state fMRI data acquired in macaque monkeys (Milham et al. 2018). We obtained tract-tracing data from a publicly available database of weighted and directed interareal connections based on a systematic large-scale anatomical investigation of the macaque cortex (Markov et al., 2013, 2014). For both types of connectivity, we assessed the geodesic distance along the cortical surface to highly connected areas. In keeping with the prior finding of a gradient extending from regions of primary cortex, we predicted that the further a cortical site is located from primary cortex, the longer its average connectivity distance. We found both the functional and structural connectivity distance to systematically vary as a function of distance from locations of primary cortex. In conjunction with our previous findings in the human cerebral cortex, the current study provides evidence for a phylogenetically conserved relationship between the spatial layout of cortical areas and connectivity.

## Materials and Methods

### Data

#### MRI data

Anatomical and functional MRI data of the macaque monkey cortex was obtained from the publicly available NKI dataset of the PRIMatE Data Exchange database (Milham et al., 2018). The Newcastle University cohort was included in the current study, consisting of 14 macaque monkeys (macaca mulatta) scanned on a primate-dedicated Vertical Bruker 4.7T scanner^1^. Analysis was restricted to 10 animals (8 males, age=8.28+/−2.33, weight=11.76+/−3.38), each of whom included two awake resting-state fMRI scans. The details of the data acquisition and experiment procedures have been described in previous studies (Baumann et al., 2011, 2015; Poirier et al., 2017; Rinne, Muers, Salo, Slater, & Petkov, 2017; Schönwiesner, Dechent, Voit, Petkov, & Krumbholz, 2015; Slater et al., 2016; Wilson et al., 2015). In brief, two 8.33-min (250 volumes) sessions of resting-state fMRI data were acquired while the animals were awake (resolution=1.2mm isotropic, TE=16 ms, TR=2000 ms), and structural T1-weighted images were acquired using MDEFT sequence (0.6mm isotropic resolution, TE=6ms, TR=750ms).

#### Tract-tracing data

Tract-tracing data of the macaque monkey cortex was obtained from the publicly available database Core-Nets (core-nets.org). Acquisition details have been described elsewhere (Markov et al., 2014, 2011). In brief, the macaque cortex was parcellated into 91 regions per hemisphere (M132 atlas). Across 28 macaque monkeys, retrograde tracers were injected into a subset of 29 regions. These tracers label neurons that project onto the injected region. Hence, we will on occasion refer to injections sites as projection sites, and regions of labelled neurons as source regions.

The current study used information on the number of labelled neurons per region for each injection. Geometric measures were taken from the cortical surface (at mid-thickness) representation of the Yerkes19 group atlas in the form of a triangulated surface mesh including the M132 parcellation and injection sites (Donahue et al., 2016).

### Data Processing

The code for data processing and analysis has been made available online ^2^.

#### MRI preprocessing

Structural and functional image preprocessing have been described in detail in Xu et al. (2017). Structural processing comprised spatial noise removal, constructing a mean image from multiple T1-weighted images, brain extraction, tissue segmentation, reconstructing native surfaces, and surface registration from native space to the Yerkes19 group template surface (Donahue et al., 2016). The surfaces were downsampled to 10,242 vertices per surface.

Functional preprocessing was conducted using a customized version of Connectome Computation System for non-human primate (NHP) data (Xu, Yang, Jiang, Xing, & Zuo, 2015). Temporal spikes were compressed and the time series were corrected for slice timing, motion, and field bias. Further steps included normalizing the 4D global mean intensity, regressing out nuisance signal (mean time series for WM and CSF masks, Friston-24 motion parameters), linear and quadratic trends, as well as band-passed (0.01 < f <0.1 Hz) filtering. The functional volumes were registered to native anatomical space and projected onto the native middle surface, where they were spatially smoothed (3 mm full width at half maximum) and downsampled to 10k surface. The preprocessed scans were then concatenated for each monkey.

The functional data was quality controlled using framewise displacement (FD) (Power, Barnes, Snyder, Schlaggar, & Petersen, 2012). We performed scan scrubbing by including time points with a mean FD below or equal to 0.2 mm. The average mean-FD across the included scans (10 animals × 2 scans) was 0.066 (SD=0.018).

#### Connectivity distance

We computed connectivity distance both from tract tracing and resting-state fMRI. While tract tracing reflects a gold-standard direct measure of cortico-cortical connections, the technique also remains limited for whole-brain studies by the number of injections. Connectivity measures obtained using resting-state fMRI, on the other hand, are inferred from correlated time courses and as such should be regarded as an indirect measure of connectivity. Its primary advantage is full-brain coverage. The conjunction of both measures allowed us to leverage the merits of each, complementing the sparse nature of tract tracing with full-brain coverage of resting-state fMRI, as well as validating the indirect measure from resting-state fMRI against the gold-standard measure of connectivity from tract tracing.

#### Functional connectivity distance

Connectivity matrices were created separately for each hemisphere of each macaque monkey. Functional connectivity between each pair of cortical nodes was quantified by correlating their time series (Pearson product-moment correlation coefficient) and were entered in a node-by-node matrix of functional connectivity.

To capture the distance of a cortical site from its functionally connected areas within the cortical layout, we computed the average geodesic distance from each node of the native surface to its functionally connected nodes. The geodesic distance describes the shortest path along the cortical surface between two points allowing a direct assessment of the implications of relative spatial positions across the cortical surface. Geodesic distance was computed using an algorithm for approximating exact geodesic distance on triangular meshes (O’Rourke, 1999) as implemented in the python surfdist and gdist packages.

The calculation of functional connectivity distance was identical to that described in our previous investigation into the human cortex (see Fig. 1 in Oligschläger et al., 2017). Connectivity was thresholded for each node by its 2% highest connectivity to determine the functionally connected nodes. Applying a node-wise threshold instead of defining an overall threshold attempts to adjust for differences in correlation strengths across nodes. For each node, the distance to 2% highest connected nodes was then averaged, yielding the overall functional connectivity distance for that node. Maps of functional connectivity distance were averaged over the monkeys and then averaged across nodes within each area of the parcellation.

**Figure 1.**
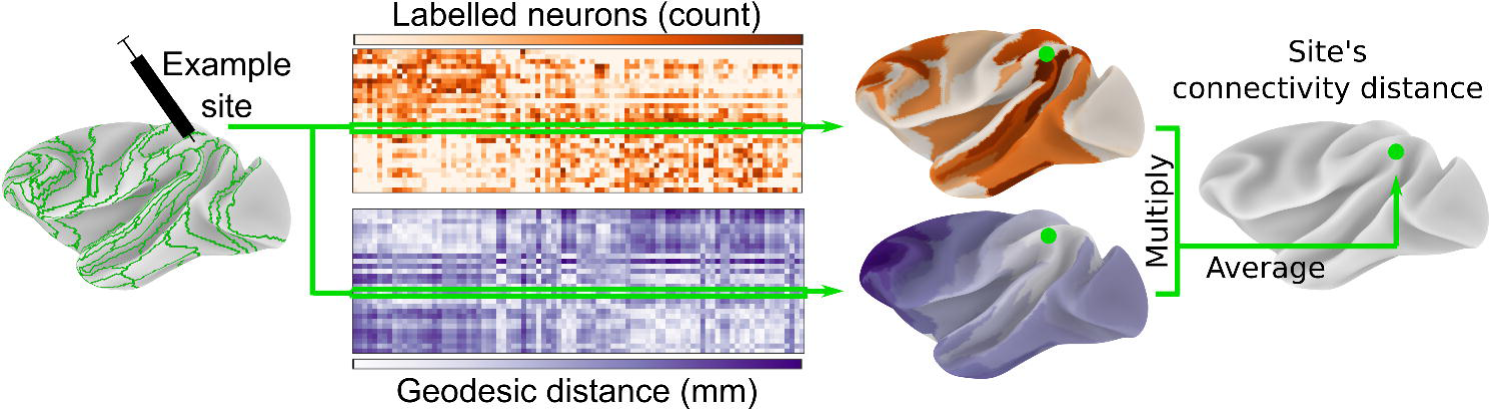
Structural connectivity distance captures the average distance of a cortical site from its source neurons. Retrograde tract-tracing data for 29 injected regions of the M132 macaque cortical atlas (green outlines) was obtained from Core-Nets (Markov et al., 2014, 2011). Geodesic distance was calculated along the cortical surface of the Yerkes19 group atlas (Donahue et al., 2016). For each injection, the connectivity distance was calculated by the injection site’s average geodesic distance from all area centroids weighted by the count of labelled neurons per area. Area centroids were used because the precise location of each labelled neuron was unknown.

#### Structural connectivity distance

To capture the distance of a cortical site from its source neurons, we calculated the geodesic distance from the injection sites to their labelled neurons. As the precise location of each labelled neuron was not reported, we instead used the centroid of the corresponding area as spatial reference for the labelled neurons. We only included labelled neurons extrinsic to respective injected area. For each injection site we then took the average geodesic distance from all its source neurons as a summary measure of its connectivity distance (Fig. 1). As source neurons within the same area shared the same spatial reference location, this is equivalent to an area’s average geodesic distance from its source areas weighted by the respective number of source neurons.

### Data analysis

We next investigated the topography of connectivity distance. Specifically, we asked whether a cortical site’s connectivity distance is related to that site’s distance from primary sensory-motor areas. Based on our hypothesis that the spatial distribution of connectivity distance follows a gradient anchored in primary cortex, we predicted that the further a cortical site is located from primary cortex, the longer its connectivity distance. For both the functional and structural connectivity distance, we tested these predictions using a general linear model (GLM), with an area’s geodesic distance from the border of the closest primary area as independent variable, and its connectivity distance as the dependent variable.

As locations of primary cortex (Fig. 2), we included primary visual (V1), primary auditory (Core), primary somatosensory (area 3), and primary motor areas (F1). We excluded olfactory and gustatory primary cortex from our analyses because they constitute chemosensory systems with distinct patterns of laminar differentiation. Unlike primary areas of visual, auditory, and somatosensory cortex, those of olfactory and gustatory cortex do not exhibit the highly differentiated structure of koniocortex (Johnson, Illig, & Behan, 2000; Pritchard, 2012; van Hartevelt & Kringelbach, 2012). While the primary areas of visual, auditory, and somatosensory systems give rise to processing hierarchies that sequentially convey information about the external environment to the least differentiated limbic structures, the chemosensory systems reflect a closer relationship to the limbic structures (Mesulam, 1998), and thus depart from the relationship hypothesized by the tethering hypothesis.

**Figure 2.**
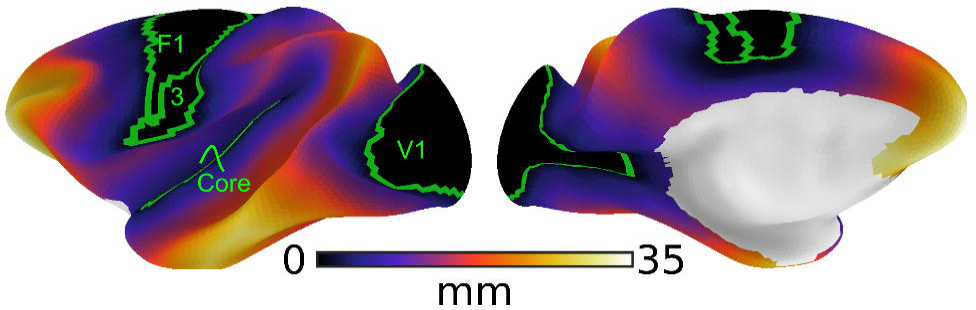
Distance from primary sensory-motor cortex. The map shows the geodesic distance from the closest border of a primary cortical region (green outline: primary visual (V1), primary auditory (Core), primary somatosensory (area 3), and primary motor areas (F1)). Geodesic distance describes the shortest path along the cortical surface and was used to assess of the implications of relative spatial positions across the cortical surface. The distance from primary cortex served as the predictor variable in GLMs assessing whether the spatial distribution of connectivity distance follows a gradient anchored in primary cortex.

For functional connectivity distance, the GLM expressed the spatial distribution of the average connectivity distance per area as a function of an area’s average geodesic distance from the closest border of primary sensory-motor areas. A major advantage of the functional connectivity here is the whole-brain coverage (compared to 29 cortical samples in the tract-tracing data).

Similarly, the spatial distribution of structural connectivity distance was modelled as a function of distance from primary cortex. Here, the geodesic distance of each injection site to the closest border of primary sensory-motor areas served as the independent variable in the GLM. Two covariates of no interest were included. First, as we only included labelled neurons extrinsic to the injected area, injections into a larger areas will have increased connectivity distance solely by virtue of the methodological setup. We thus corrected for region size as a confounding factor by including the number of nodes in the injected region as a covariate in the GLM. Second, some areas are positioned within the cortical geometry to have overall longer distances to the rest of the cortex than geometrically more central areas. The effect of position within the overall geometry of the cortex was thus measured as the average distance of an injection site from all cortical nodes and added as a covariate to the GLM.

In addition to the hypothesis-driven analysis describing the spatial distribution of connectivity distance with respect to locations of primary cortex, we conducted a search procedure to check for the possibility that the distance from a different set of regions might account better for the spatial distribution of structural connectivity distance. To this end, we randomly sampled cortical locations 100,000 times and repeated the above-described GLM using these locations as anchor points instead. To identify good reference locations, we calculated each node’s mean variance explained across all GLMs that included this node in their random set of anchors. The resulting map of average variance explained provides a hypothesis-free estimate of locations that anchor the spatial distribution of connectivity distance.

However, probing the search space of all possible sets of reference nodes by random sampling poses a huge combinatorial problem which we addressed and replaced in a second and separate analysis that included optimization. Starting with a random set of nodes (size between 3 and 5), these nodes were moved in steps of 1 mm towards the direction of higher variance explained until a stable set of locations was found (i.e. where further moving by 1 mm did not result in higher variance explained). This search was repeated 1000 times, each time starting with a new random set of nodes. For each node, we counted the number of times it was included in the final and stable set of reference nodes. The resulting map of counts distinguishes between positive (the further from reference location, the longer the connectivity distance) and negative (the further from reference location, the shorter the connectivity distance) relationships.

## Results

### Functional connectivity distance follows a gradient anchored in primary cortex

We investigated the spatial distribution of connectivity distance based on resting-state functional connectivity in relation to locations of primary cortex. Connectivity distance was shortest in sensory-motor regions and peaked in higher-order association regions in lateral and medial frontoparietal cortex (Fig. 3A). Information on each area’s connectivity distance can be found in Online Resource Fig. 1.

**Figure 3.**
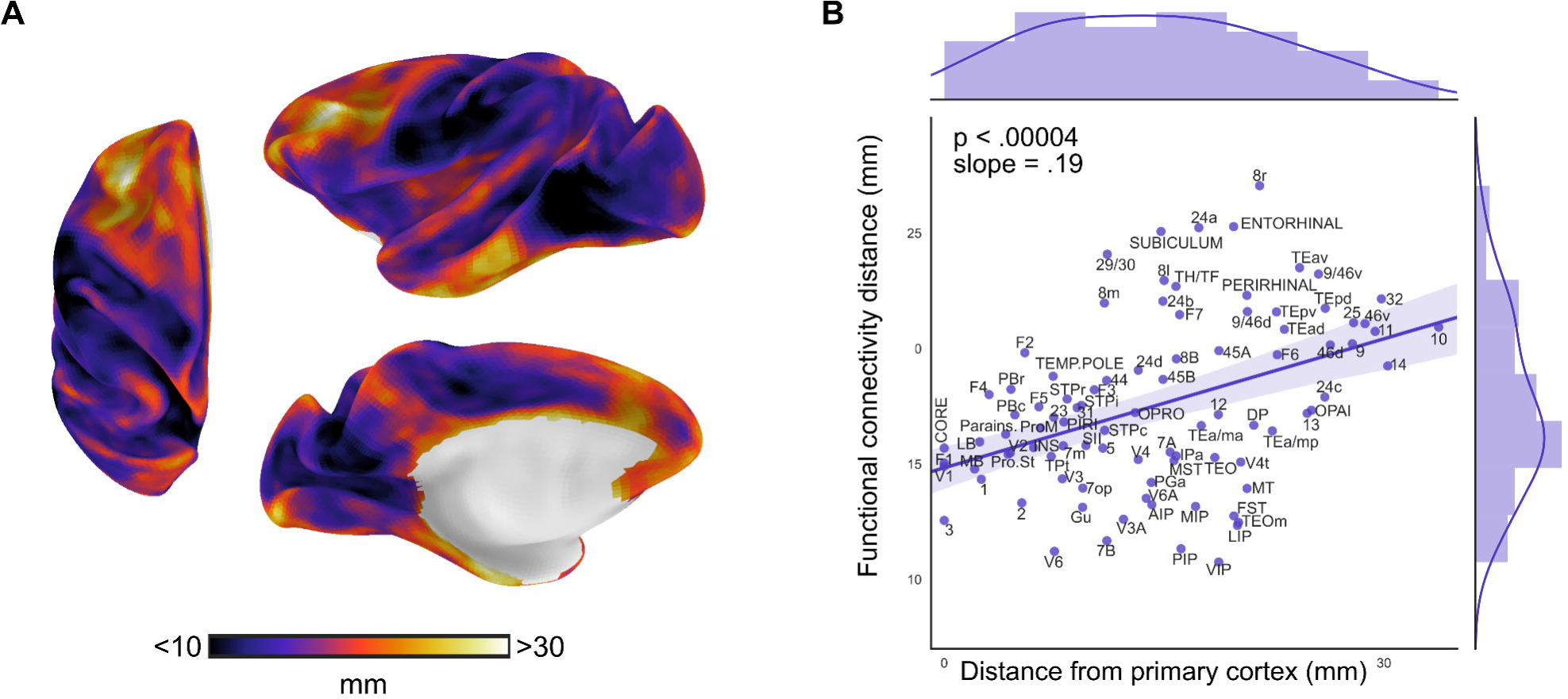
The spatial distribution of functional connectivity distance (A) was investigated with relation to locations of primary cortex. (B) Each area’s mean functional connectivity distance is plotted against its mean distance from primary cortex. Functional connectivity distance systematically varied across the cortex as a function of distance from primary cortex demonstrated by the significant relationship between the map of distance from primary cortex (shown in Fig. 2) and connectivity distance (A) (F_1,89_=18.8, p<.00004, R^2^=.17; t_90_=4.3, p<.00004, beta=.19, 95% CI=[.10, .27]). These findings show that the further a node was located from primary cortex, the longer its functional connectivity distance.

We found functional connectivity distance to systematically vary as function of distance from primary cortex. A GLM with distance from primary cortex as independent variable and connectivity distance as dependent variable (F_1,89_=18.8, p<.00004, R^2^=.17) showed that the distance from primary regions significantly accounted for the spatial distribution of functional connectivity distance (t_90_=4.3, p<.00004, beta=.19, 95% CI=[.10, .27]). Specifically, the further a node was located from primary cortex, the longer its functional connectivity distance (Fig. 3B).

By an arbitrary threshold of studentized residuals larger than 2, the functional connectivity distance of areas VIP, 8r, as well as cortical limbic regions 24a, 29/30, subiculum, and entorhinal cortex were found to deviate from the overall pattern. While VIP is at medium distance from primary cortex, it had among the shortest functional connectivity distance. Other intraparietal areas (PIP, LIP, MIP) shared this property but deviated less. In contrast, area 8r and the cortical limbic regions had much longer functional connectivity distance than predicted by their distance from primary cortex. Information on each area’s deviation from the overall relationship is presented in Online Resource Fig. 2.

### Structural connectivity distance follows a gradient anchored in primary cortex

The spatial distribution of structural connectivity distance was also inversely related to distance from primary cortex. While the functional connectivity analysis was calculated across the entire cortex, the number of injection sites restricted our analysis to the spatial distribution of injection sites with respect to primary cortical areas.

Structural connectivity distance was shortest in unimodal cortex and highest in frontoparietal regions of higher-order association cortex (Fig. 4A), and followed a gradient extending from primary cortical areas. The GLM (F_3,25_=10.01, p<.0002, R^2^=.55) revealed a significant relationship between structural connectivity distance and distance from primary cortex. Connectivity distance systematically increased with distance from primary cortex (t_26_=2.12, p<.05, beta=.18, 95% CI=[.005, .35]). Thus, the further an injection site was located from primary cortex, the longer its structural connectivity distance (Fig. 4B). Information on each injection site’s connectivity distance can be found in Online Resource Fig. 1.

**Figure 4.**
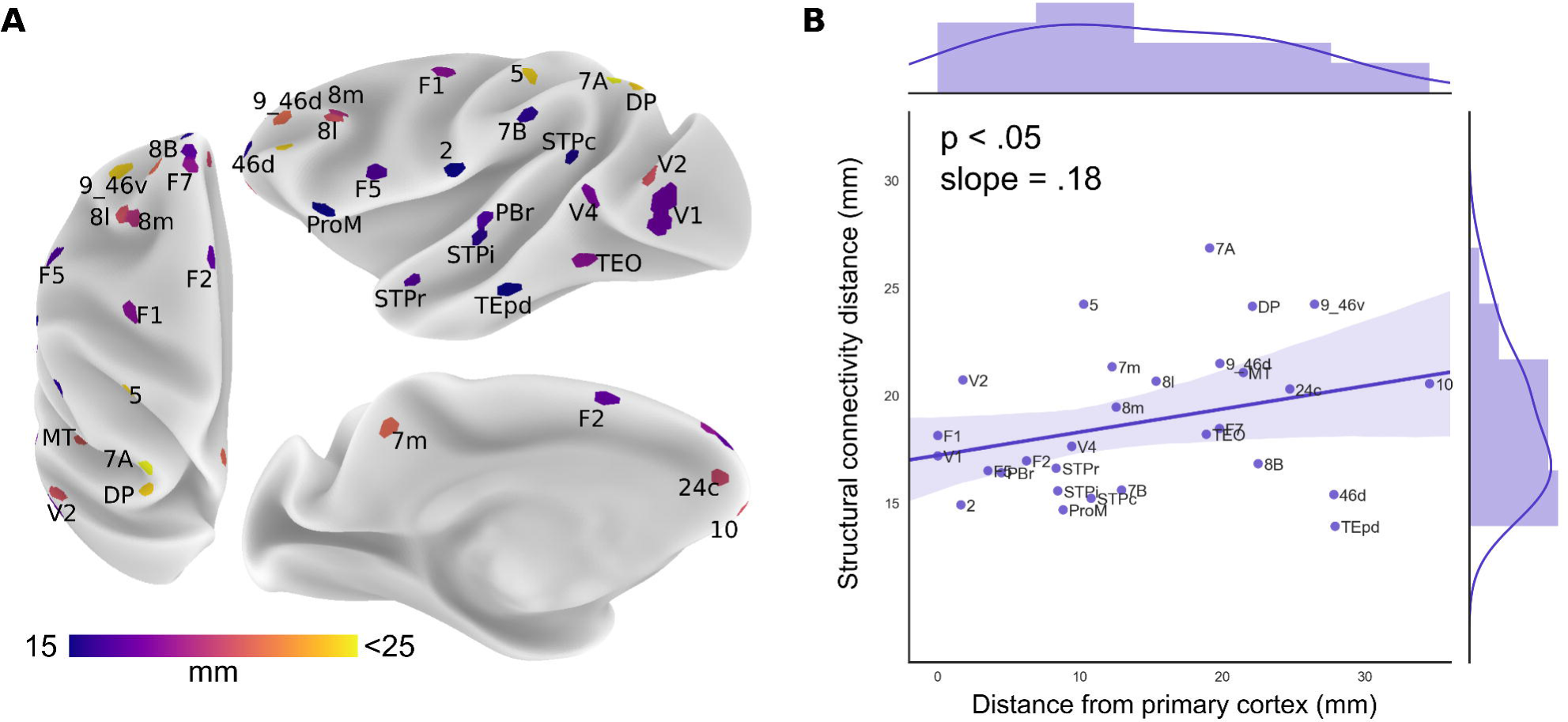
The spatial distribution of structural connectivity distance (A) was investigated with relation to locations of primary cortex. Structural connectivity distance was corrected for confounding factors of region size and location. (B) Structural connectivity distance systematically varied across the cortex as a function of distance from primary cortex demonstrated by the significant relationship between the map of distance from primary cortex (shown in Fig. 2) and connectivity distance (A) F_3,25_=10.01, p<.0002, R =.55; t_26_=2.12, p<.05, beta=.18, 95% CI=[.005, .35]).

The structural connectivity distance of areas TEpd and 7A deviated from the overall pattern (by a studentized residual > 2). While TEpd is far from primary cortex, it had among the shortest structural connectivity distances. In contrast, area 7A had the longest structural connectivity distance despite not being the furthest away from primary cortex. Both areas also differed substantially in their structural and functional connectivity distance. In the functional analysis, these areas follow the trend with respect to location of primary cortex more closely. Further information on deviations from the overall relationship are presented in Online Resource Fig. 2.

### Spatial distribution of structural connectivity distance (Search procedure)

In addition to the hypothesis-driven analysis describing the spatial distribution of connectivity distance with respect to locations of primary cortex, we conducted a search procedure to check for the possibility that a different set of regions might account better for the spatial distribution of the structural connectivity distance. A map of the average variance explained across nodes was obtained from 100,000 GLMs. Instead of distance from primary cortex, these GLMs included the distance from random locations as independent variable. The map (Fig. 5) identifies regions for which the distance from them either positively (red) or negatively (blue) account for the distribution of structural connectivity distance. These maps offer an estimate of the whole-brain topography of structural connectivity distance based on the subset of known injection sites.

**Figure 5.**
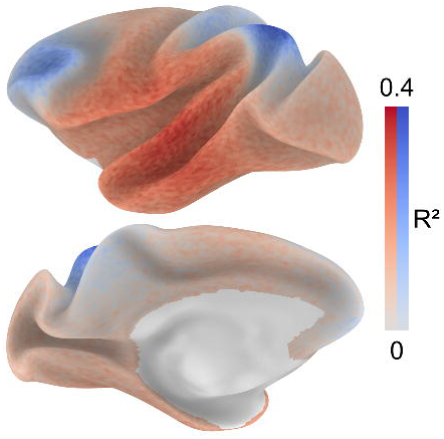
Hypothesis-free search for anchor regions. To estimate locations that anchor the distribution of structural connectivity distance, 100,000 GLMs were repeated using distance from random sets of locations as the predictor variable. For each surface node, we calculated the average variance explained across the GLMs that included distance from a given node as the predictor. The map identifies regions from which the distance either positively (red) or negatively (blue) account for the distribution of structural connectivity distance. These maps offer an estimate of the whole-brain topography of structural connectivity distance based on the subset of known injection sites.

A comparison of functional connectivity distance with structural connectivity distance (Online Resource Fig. 3) and with estimated whole-brain structural connectivity distance (Online Resource Fig. 4) did not reveal a similarity across modalities. While the topography is partly consistent with that of the available whole-brain map of functional connectivity distance, differences pertain to regions of occipitoparietal and temporal cortex. Specifically, areas V2, MT, DP, 7A, 5, 7m, and area TEpd deviated the most from an ideal linear fit between functional and structural connectivity distance (Online Resource Fig. 3).

A separate search optimization procedure confirmed these findings. 1000 random locations were moved in 1-mm steps along the cortical sheet until a stable set of locations was found that best predicted the distribution of connectivity distance. We counted the number of times each node was included in the stable set of reference nodes (Online Resource Fig. 5). Peak regions confirm with the reported regions from the initial search procedure.

## Discussion

The current study demonstrates a relationship between the spatial layout and connectivity of the macaque monkey cortex. The conjunction of tract-tracing and resting-state functional connectivity data allowed us to leverage the merits of each, complementing the sparse nature of tract tracing with full-brain coverage of resting-state fMRI, as well as validating the indirect measure from resting-state fMRI against the gold-standard measure of connectivity from tract-tracing. Characterizing a region’s connectivity distance – measured as a region’s average distance to connected areas - we found that the spatial distribution of connectivity distance follows a gradient that is anchored in primary cortex. While a direct similarity between structural and functional connectivity distance is only partly present, both measures follow this gradient. Specifically, the further a region is located from primary cortex, the stronger its connections to distant portions of the cortex. This topographic pattern is consistent with our previous findings in the human cortex (Oligschläger et al., 2017) offering an evolutionary perspective on brain organization.

The topography of connectivity distance in the macaque monkey cortex presented here coincides with that of the human cortex (Oligschläger et al., 2017). While deviations were observed in the lateral temporal and cingulate cortex across species, the overall similar topography of connectivity distance between the macaque and human cortex suggests an organizational principle of cortical connectivity. This correspondence opens up the possibility of a phylogenetically conserved relationship between connectivity and the spatial layout in the cortex. Nevertheless, given the substantial phylogenetic distance between humans and macaque monkeys, future studies are needed to investigate this relationship in across a broader spectrum of primate as well as other mammalian species.

An evolutionary perspective on cortical organization has been proposed by Buckner and Krienen (2013) who attribute the distribution of connectivity distance to the cortical expansion that occurred during primate phylogeny. Indeed, the topography of connectivity distance in both the macaque and human cortex closely concur with patterns of phylogenetic expansion (Chaplin, Yu, Soares, Gattass, & Rosa, 2013; Fjell et al., 2015; Hill et al., 2010). According to the ‘tethering hypothesis’ (Buckner & Krienen, 2013), cortical expansion during primate evolution has caused increasingly larger amounts of cortex to lie distant from primary cortex. Untethered from developmental constraints that differentiate primary cortex, these regions form unique characteristics of association cortex, such as their long-range connections.

Alongside the overall agreement between connectivity distance in the macaque and human cortex, we found a notable difference in the lateral temporal cortex of the macaque. While this region was among those with the longest functional connectivity distance in both the human and macaque monkey, it had comparatively short structural connectivity distance in the macaque. This set of observations does not afford to differentiate a true species difference from a method-related difference. Assuming a true species difference, however, this would be consistent with previously reported differences between the human and macaque neuroanatomy and connectivity. The human temporal lobe is disproportionately larger (Rilling & Seligman, 2002) and marked by strongly modified projection patterns of the arcuate fasciculus (Rilling et al., 2008).

Further differences between the topography of macaque and human connectivity distance include posterior cingulate cortex, rostral anterior cingulate cortex (ACC), and posterior insula. In the human cortex, these regions deviate from the overall pattern and show relatively short connectivity distance compared to surrounding regions (Oligschläger et al., 2017). These deviations were less evident in the macaque. However, in the macaque too, the insula is marked by relatively short connectivity distance similar to sensory-motor regions, and the ACC exhibits a faint decrease in connectivity distance.

Our findings join a body of research into organizing principles that govern cortical connectivity of the mammalian brain. The distance between cortical sites is a major determinant of cortical connectivity, with connectivity strength between regions systematically decreasing with the distance between them (Betzel et al., 2016; Ercsey-Ravasz et al., 2013; Roberts et al., 2016). The findings presented here suggest that connectivity distance is modulated by a region’s offset from primary cortex. Specifically, the rate by which connectivity strength decreases with the distance between regions might be modulated by how far they are located from primary cortex. As the connectivity strength has been shown to follow an exponential decay with distance (Ercsey-Ravasz et al., 2013), we expect this decay to be steepest in primary sensory-motor areas, becoming progressively less steep with distance from these areas.

The current findings provide spatial context in which organizing principles operate and might prove useful for integrating these principles with potential developmental mechanisms that give rise to them in the first place. A promising line of research models spatiotemporal dynamics during cortical ontogeny to reproduce major characteristics of connectivity. For example, assuming axons connect with the first neuron they encounter, models of random axonal outgrowth successfully reproduced patterns of decreasing connectivity with distance (Kaiser 2009). In a serial version of this model, neurons that started axon growth early formed longer connections than later ones (Lim and Kaiser 2015). This is of special relevance for the current findings given that neurons destined for association areas migrate earlier during cortical ontogeny than primary cortical areas (Figure 2 in Rakic, 2002).

Postnatal cortical maturation follows an opposite spatiotemporal pattern. While sensory-motor regions mature early during childhood, it is association cortex that fully matures last (Casey, Tottenham, Liston, & Durston, 2005; Chomiak & Hu, 2017; Toga, Thompson, & Sowell, 2006). This spatiotemporal trajectory characterizes the maturation of structural properties such as gray matter density (Gogtay et al., 2004), thickness (Sowell et al., 2004), and surface expansion (Hill et al., 2010), as well as synaptic density (Huttenlocher, 1990) and myelination (Flechsig, 1920). When association cortex reaches maturity, it is marked by a more elaborate and complex microstructure than sensory-motor cortex. For example, pyramidal neurons in association cortex are larger (Elston, Tweedale, & Rosa, 1999) and possess both a larger number as well as longer dendrites (Elston, Benavides-Piccione, & DeFelipe, 2001). It has been proposed that these structural properties allow association regions to integrate larger numbers of inputs compared to primary-sensory motor regions (Elston, 2002). The current findings provide further support for this proposal by indicating that association cortex integrates input from more widespread cortical regions.

Thalamocortical connectivity, established during cortical ontogeny, is another candidate mechanism that might define the observed topography of connectivity distance. While unimodal sensory-motor regions receive input from their specific thalamic nuclei, distributed regions of association cortex share thalamic input from the medial pulvinar nucleus (Goldman-Rakic, 1988). For unimodal regions, it has been shown that thalamic input determines activity-dependent differentiation of sensory hierarchies (Chou et al., 2013; Katz & Shatz, 1996; Lokmane et al., 2013). An interesting possibility is to consider a similar mechanism for the distributed regions of association cortex. These, too, might interconnect in a similar activity-dependent manner via their common thalamic input (Buckner & Krienen, 2013; Shipp, 2003). In conjunction with such models, an understanding of basic organizational principles of cortical connectivity can inspire future investigations that study these principles as an emergent property of spatiotemporal processes during cortical ontogeny.

The emergence of areas over the course of cortical phylogenesis might further account for the observed topography of connectivity distance. A connectivity ‘blueprint’ may be established during cortical evolution such that regions predominantly connect with those already in existence when they evolve. Hence, a general tendency might be that connections are established mainly to regions that are phylogenetically older, but not vice versa. In that sense, the phylogenetically more recent regions of association cortex would be expected to connect both to sensory-motor cortex and among themselves, while sensory-motor areas connect primarily within their cortical hierarchies. Such scenario would imply a dissociation between sensory-motor and association cortex with respect to afferent and efferent connectivity distance. While association cortex would be marked by both long efferents and afferents, sensory-motor cortex would be marked by long afferents and short efferents. As the sequence of emerging areas during cortical evolution is a controversial question (cf. Sanides, 1969), future studies that incorporate measures of the directionality of connections could help shed further light on such questions.

From a functional perspective, connectivity distance might index a region’s hierarchical level and integrative capacity of information processing. For example, prior findings from Wagstyl and colleagues (2015) demonstrate that geodesic distance from primary cortical areas traces structural hierarchies in the macaque monkey cortex. Assuming that the level of global connectedness of a region contributes to its capacity of information integration, connectivity distance may be a structural feature that affords hierarchies of information integration in the cortex.

To conclude, we reported on a phylogenetically conserved relationship between cortical layout and connectivity of the primate cortex. As in the human cortex (Oligschläger et al., 2017), we revealed a gradient of connectivity distance extending between primary and frontoparietal regions. The further a region is located from primary sensory-motor cortex, the stronger its connections to distant portions of the cortex. Together, these findings highlight the utility of cross-species comparative studies for addressing the phylogeny of cortical organization.

## Compliance with Ethical Standards

The authors have no conflicts of interest to declare. All data used in the current study are openly available and were previously acquired in accordance with ethical standards of respective institutions.

## Footnotes

1 http://fcon_1000.projects.nitrc.org/indi/PRIME/newcastle.html

2 https://github.com/soligschlager/distconnect_macaque/tree/master

